# Maximization of non-nitrogenous metabolite production in *E. coli* using population systems biology

**DOI:** 10.1101/2019.12.27.888545

**Authors:** Sreenath Rajagopal, Arindam Ghatak, Debdatto Mookherjee, Rothangmawi Victoria Hmar, Anirudh P. Shanbhag, Nainesh Katagihallimath, Janani Venkataraman, KS Ramanujan, Santanu Datta

## Abstract

The *E. coli* metabolome is an interconnected set of enzymes that has measurable kinetic parameters ascribed for the production of most of its metabolites. Flux Balance Analysis (FBA) or Ordinary Differential Equation (ODE) models are used to increase product yield using defined media. However, they either give a range (FBA) or exact amount (ODE) of metabolite yield which isn’t true as the transcriptome diversity of individual cells isn’t considered. We formulate the metabolic-behaviour of individual cells by using a POpulation SYstems-Biology ALgorithm (POSYBAL) which predicts multiple-gene knockouts for increasing industrially relevant metabolites. We validate this prediction for producing isobutanol (Heterogenous metabolite) and shikimate (Homogenous metabolite) where, the product-yield was increased by 40 times (~2000 ppm) and 42 times (~3000 ppm) respectively. Also, we introduce a nitrogen-swap in standard media to its low-nitrogen counterpart during post-growth phase to redistribute flux towards non-nitrogenous pathways for increasing overall product-yield. Further, our model shows growth-phase diversity in bacterial population even under normal glucose-uptake, portraying a real-world scenario of diverse and robust environment thus, making it evolutionarily favourable to threats such as anti-bacterial attack.

## Introduction

Naturally growing bacteria have evolved to survive in the “occasionally famine and rarely feast” conditions for biomass. In contrast, the nutritional factors are made optimal in laboratories to return higher biomass, subsequently increasing the product yield. Inherent in this optimizing process is often an opposite pull of flux of either maximizing the biomass which is a near precise stoichiometric summation of multiple essential metabolites or the specific maximization of any particular metabolite. If these ‘nutritional conditions’ can be exploited, we can have a ‘minimalistic’ system to carry out reactions of commercially valuable metabolites with low-cost inputs. To this end, we need to construct an in-silico model and validate its formulation by precise experimental operations.

One of the significant drivers for constructing in-silico models for different micro-organisms in the field of metabolic engineering is to forecast the genetic changes (gene deletions and over expressions) required in enhancing the levels of a specific metabolite of the interest. There are two types of platform that are commonly used to simulate a bacterial cell. It is either based on linear Flux Balanced analysis (FBA) platform (Burmeister M, 2007, Latendresse *et al*, 2012, Orth *et al*, 2010, Raman *et al*, 2008) or dependent on ordinary differential equations (ODE) where kinetic parameters (Km and Vmax) of enzymes participating in the particular reaction type (single substrate, multi-substrate, ping pong, ternary complex etc.) are used (Bowden *et al*, 1999, Kim *et al*, 2018, Mannan *et al*, 2015, Suresh *et al*, 2010). The popular FBA platform generally incorporates the development of a stoichiometric model with genome-level annotations of pathways that maps conversion of xi moles of substrates to xj moles of product. The simulations thereof are used to predict knockouts of non-essential genes, in specific pathways (identified during the simulation studies) which would help in blocking the formation of particular metabolites and allow the cells in redirecting the carbon feed into the production of the metabolite of interest.

All types of in-silico model, whether based on the mathematical framework of FBA or ODE format tacitly assume that all the cells are in an identical metabolic state. However, bacterial population are generally asynchronous in their growth and their reaction to a stimulus (positive or negative) at the individual cellular level are not unique and can only be represented by a diverse array of responses. In a population of bacterial cells under a standard nutrient condition, some proliferate while some are stunted. Similarly, upon exposure to an antibacterial compound over a period of time, say about 90-99 % of the population are affected, while the remaining 1-10% remain unaltered due to the robustness in the system. For eg. the heterogenous expression of araBAD promoter in the presence of limited arabinose quantities in *Mycobacterium tuberculosis* shows variation in individuals of a population (Yang *et al*, 2011). Additionally, as summarized in a recent review (Leyberger *et al*, 2019) it is a truism that in a population of bacterial cells, the gene expression levels are highly variable in lower nutrient concentrations.

Traditional modelling assumes every bacterium to have equal exposure to the participating nutrients. Such a ‘socialistic’ approach is often invalidated in a natural environment where intra-species competition persists leading to crony capitalism, among bacteria where nutrient availability is the ‘currency’ for driving a reliable binary of ‘haves’ and ‘have-nots’. In essence, the collection of cells in a group with the same genetic makeup (isogenic) have different expression profile. Hence, there is an urgent need to develop a population model where each cell has a unique metabolic signature.

For understanding the optimal flux of pathways in a population, we can use a similar approach. Several industrially relevant metabolites are devoid of nitrogen. With glucose as the sole carbon source, the anabolism of nitrogen-containing metabolites such as nucleotides and amino acids, are further downstream compared to the C, H, O based metabolites. Hence, it is easier to use the POSYBAL platform owing to the requirement of lesser dimension of matrices and hence lower computing power. We describe the mathematical foundations of a population model extending the FBA architecture and validate its predictive power by constructing strains of *E. coli* with minimal genetic modifications that produce non-nitrogenous metabolites such as Isobutanol and Shikimate at multigram levels.

Isobutanol is produced by altering the pathway from the formation of branched-chain amino acids (BCAA), whereas shikimate is an intermediate metabolite of the aromatic amino acid pathway (AAA) respectively. Isobutanol is an important industrial solvent which has higher calorific value than ethanol making it an ideal fuel substitute. It requires no infrastructure modifications for transport, and unlike ethanol, it is not hygroscopic and is non-corrosive to motor engines. The generation of CO2 instead of SO2 or CO makes it a clean fuel compared to the fuels derived from petroleum. On the other hand, shikimate is a high-value industrial precursor for producing herbicide such as Glyphosate and the antiviral compound Tamiflu. (Knaggs *et al*, 2002).

Both Isobutanol and shikimate are produced by designed alteration of the BCAA (Atsumi *et al*, 2008) and AAA pathways respectively. Fig 2 illustrates the metabolic pathways required for their production. The unique feature of all the intermediates/metabolites that are formed from glucose to isobutanol and shikimate (Fig 2) is the absence of nitrogen in the metabolite composition. Hence, after producing the biomass in a rich media we generated the metabolite by using media containing either zero or limited amount of nitrogen. This was done to force the flux towards the production of isobutanol and shikimate. We term this shift of media as ‘Nitrogen Swap’ (N-swap) since it is literally the swapping of nitrogen to obtain more product yield. Of course, nitrogen is an essential element for cellular growth and is a part of proteins and DNA/RNA. Hence under nitrogen depleted condition, the growth of the cell is halted. However, since carbon, hydrogen and oxygen are present, the metabolite flux does proceed through Nitrogen independent pathway.

Additionally, we use BL21 strain, a B-Strain of *E. coli* the strain has a valine-feedback independent acetolactate synthase (ilvG). This, in addition to the heterologously expressed ketoisovalerate decarboxylase (KivD), is better suited compared to the K12 (BW25113) strain for isobutanol production. Similarly, BL21 gave better Shikimate yield than BW25113, albeit for unknown reasons (Supplementary Fig S1).

An additional advantage of N-swap in case of Isobutanol production is that under anaerobic growth conditions nitrate formation inactivates the enzyme Dihydroxy-acid dehydratase (IlvD). Dinitrosyl iron complex bound IlvD is an inactive enzyme complex making the bacteria a BCAA auxotroph effectively halting isobutanol production. However, under limiting or no nitrogen in N-swap condition, the NO formation is effectively stunted, which helps in keeping the flux active through the BCAA pathway. This complex is activated under the aerobic condition without the formation of a new enzyme (Ren *et al*, 2008). We demonstrate a scheme which considers the minimal amount of input with nominal gene manipulations necessary for the production of isobutanol and shikimate in the BL21 *E. coli* strain.

## Methods and protocols

### Population system biology algorithm (POSYBAL)

In an FBA model, a matrix (M) of size y * z is generated with stoichiometric metabolic reactions. The compounds participating in the metabolic reactions marks the rows of the matrix (y unique compounds participating in the system) and the columns represents the reactions (z overall reactions in the system). The matrix is constructed using the stoichiometric coefficients of each of the respective metabolites participating in a reaction, traversing left to right one column at a time. A positive coefficient is considered when a metabolite is produced, and negative coefficient for consumption. The number of metabolites participating in each reaction are limited, which leads to the development of a sparse matrix. The flux through each of the reactions is an unknown and is represented by a vector k of length z. Under steady state condition the rate of change of concentration of each of the metabolites is 0. Thus, any solution of k that satisfies the equation M*k = 0 is part of the null space of M (Fig 1D). The system of equations obtained by steady state mass balance is such that the total number of equations is y (equal to the No. of metabolites) and unknown variable is the flux of each reactions participating in the system z (No. of reactions). Given any large metabolic model, the number of reactions is always greater than the metabolites. When the number of unknowns is greater than the equations, we can identify many plausible values for the unknowns that are solutions of the system of linear equations. Such system of linear equations is known to be underdetermined. With the help of constrains, we can define the range for the solution space. The constraints are designed using the composition of the media used for the growth of the bacteria. In conventional FBA approach an objective function is designed that works for maximisation or minimisation of a particular reaction, giving the contribution of each of the reactions in the system towards the maximum or minimum flux of the target reaction. Individually, most enzymes intrinsically, are reversible but the forward reaction is significantly higher than the reverse reaction.

**Figure 1:**
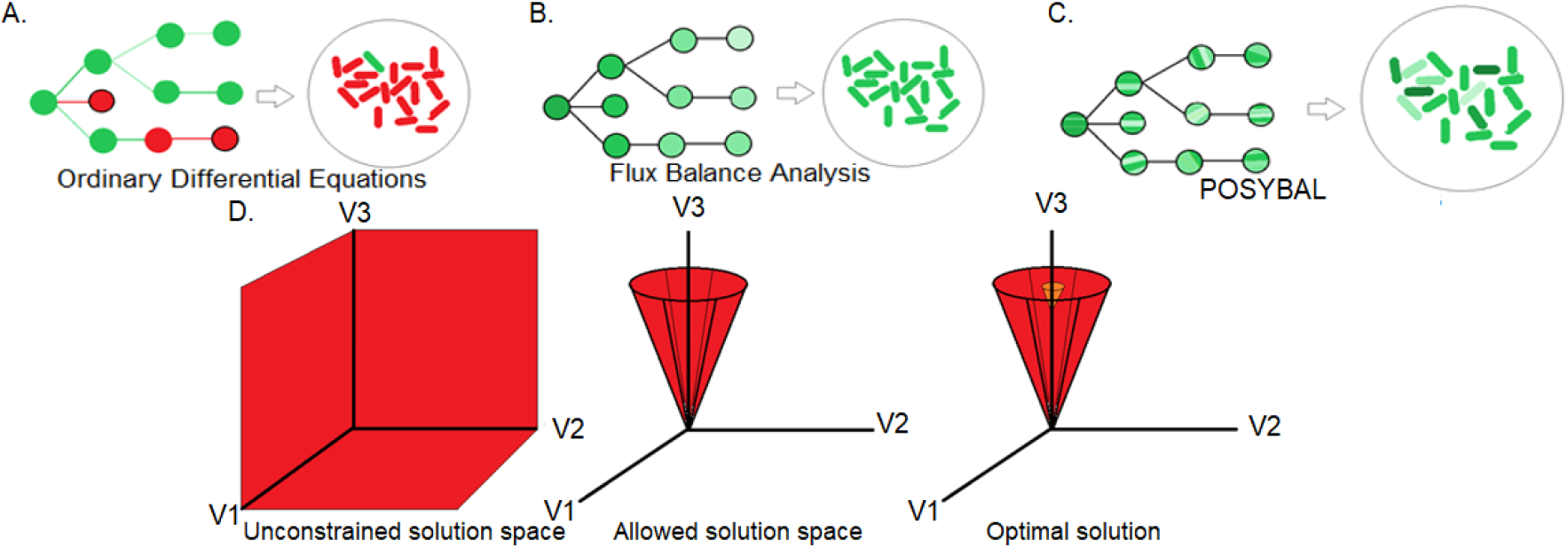
Depiction of ODE, FBA and POSYBAL models. A. The ODE model takes in the kinetic parameters and gives solutions based on those parameters. The solution is singular (green) however, gene lethality can also be found (red) for a given knockout. B. The FBA model uses stoichiometric constraints and gives a single optimal solution for the entire population. The essentiality/lethality is often found through literature or experimental evidence. C. The POSYBAL model may not predict essentiality but it considers multiple solutions for each stochiometric constraint and gives multiple combination of flux the best knockouts are chosen through the population. D. Concept behind constraint-based modelling where a solution space is limited given the ‘dimensions’ (red) such as stochiometric flux, metabolite produced and biomass. Off this optimal solution (orange) is chosen for further testing.

For our model, we have made it unidirectional for computational convenience. Fig 1D shows the reduction of solution space (red) where eventually an optimal solution (Orange) is obtained in traditional FBA by maximizing or minimizing the target reactions (Orth *et al*, 2010). Instead, in POSYBAL platform, the samples are fetched from the entire constrained solution space using the random walk methodology of Markov Chain Monte Carlo (MCMC) method. The high dimensional nature of the system of equations as found in metabolic network models makes the MCMC based approaches to be efficient for picking samples from the Null Space. The inequality constrains defines the boundary in the feasible region (hyperplane). All the points on one side of the hyperplane satisfies the inequality and are considered to be a feasible solution and the points that falls on the other side of the inequality forms infeasible solutions. The solutions that are considered for the analysis are those that satisfies all the inequalities and are found to be present in the constrained feasible region. A point is selected within the solution space where all the inequalities are satisfied acting as a seed, the sampling of the new point is done from a normal distribution with mean zero and a fixed standard deviation (jump length). The new points selected randomly depends only on the previous point. The new point is accepted if it satisfies the inequality else, we go back to the previous point and try again. The xsample () function available in the R, part of the LIM package allows us to implement this algorithm (Meersche *et al*, 2009)

We have created the shikimate model from IJO1366 (Orth *et al*,2011) with the addition of a reaction corresponding to the export of the shikimate metabolite. In the absence of a transport flux the system considers the shikimate metabolite to be unused when reactions contributing to the utilization of the shikimate are knocked out, bringing down the shikimate flux to 0. This prevent us from arriving at knockouts that can maximize shikimate. In a different approach one need not optimize the system of linear equation for a given objective function but try and see the solution distribution over a very large set of solutions within some boundary. This method tries to work around the optimization problem where it is inherently assumed that the system has some definitive but unknown intelligence to work towards (turn on genes expressions suitably) to attain maximization or minimization of the unique function. The modified approach that is computationally very intensive looks to setup a bounding set of minimal constraints so that all solutions that are possible to lie within a bound and then try and generate sample solutions from the entire solution space obtaining a matrix of multiple solutions (POSYBAL) using an algorithm based on Markov chain (Van Oevelen *et al*, 2009) and this data comprises of all the possible solution (ways) in which the organism can act (population behaviour) corresponding to a given condition. The POSYBAL data comprising of 100000 samples was developed (See supplementary methods 2 and 3 for scripting and output information).

Once the population results are obtained (i.e. matrix of multiple solutions), the system is filtered for the samples from the solution space with maximum production of the target metabolite. The maximum flux is identified and all the samples which constitutes ~90% and above of the maximum flux of target metabolite are filtered out. The maximum flux of all individual reaction is also computed. The next step is to identify the reaction fluxes that contributes least in the population level towards the production of target metabolite. We achieve this by adding a filter to sample out the reaction fluxes that run minimal ~10% of the maximum flux for that respective reaction amongst the samples with maximum yield of target flux. These indicate knockouts for validating the platform in vitro. program using R programming (R development core team, 2013) and python (Python Software Foundation, https://www.python.org) is built to filter the possible knockouts from a population. The GitHub links for the POSYBAL platform for shikimate and isobutanol synthesis is as follows

https://github.com/sreenathraiagopal/shikimatemodel_codes.git

https://github.com/sreenathrajagopal/Isobutanolmodel_codes.git

### Generation of predicted *E. coli* knockouts

Both the triple knockouts i.e. ΔackA:ΔadhE:ΔldhA and ΔaroA:ΔaroK:ΔaroL were generated using P1 transduction method (See supplementary methods). ΔackA (BW strain) was used the donor and the BL21 strain was used as the recipient. Subsequently, ΔadhE and ΔldhA were used to knockout the respective genes in BL21. Similarly, BW ΔackA strain form in-house KIEO library of *E. coli* knockouts which have a kanamycin cassette in place of the knocked-out gene as a marker. After successful transduction, BL21 ΔackA strain is made with Kanamycin cassette replacing the knocked-out gene. This is confirmed by colony PCR. For shikimate production ΔaroA from KIEO collection was used as the recipient strain. To further knockout the genes, the kanamycin marker is first flipped out by using the λ-red recombinase method. pCP20 plasmid is first transformed into the desired knockout and plated on LB-amp (30ug/ml) plate and incubated overnight. The colonies are then grown in Luria broth until they reached 0.6 OD_600_. Then, the culture is incubated at 37°C for one hour followed by incubation of 43°C for four hours. It is then, plated on plain LB media, media containing ampicillin and media containing kanamycin respectively. Growth of the culture on LB plate and no growth on media having either ampicillin or kanamycin confirmed the absence of kanamycin cassette.

### Protocol for isobutanol and shikimate production with various knock-outs and media swap

Initially, the conformation of in-silico simulations for nitrogen modulation was done in shake flasks before proceeding towards biotransformation in bioreactors. The desired knockouts and wild-type strains are then transformed with pUC57a kivD plasmid and plated on Luria agar plates with ampicillin (100ug/ml). A single transformant is then inoculated in 5ml of LB media with ampicillin (100ug/ml) and grown overnight. The pUC57-kivD plasmid, synthesized by Genscript, is engineered without an operator site and with a constitutive promoter therefore making the induction step void. For shikimate the cells (or knockouts) are grown overnight in LB media.

### Shake-flask bioconversion experiments

To conduct the shake-flask experiments for both shikimate and isobutanol growing the starter cultures in LB media, they are transferred to fresh Lysogeny broth and grown until 2.0 OD 600 (Secondary culture). Then the cells are centrifuged at 4000g and transferred into nitrogen deficient media. In case of Isobutanol, the knockouts are transferred to M9 media with varying percentage of Ammonium Sulphate (Nitrogen source) composition with 3.6% glucose (Carbon source) and ampicillin (100ug/ml) for sustaining the pUC57-kiVD plasmid. For shikimate production, the cultures are transferred into M9 media with varying percentage of LB with 1.6% glucose.

### Protocol for bioreactor

To get higher product yield the cells were grown in 500ml Bioreactor (Applikon miniBio). For producing isobutanol and shikimate the cells were grown up to ~ 6.0-6.5 OD_600_ in 20% dissolved oxygen (DO) at 200 rpm impeller speed. PEG400 was added as an anti-foaming agent. To create microaerophilic conditions the DO is reduced to 2.5% and impeller speed is reduced to 50 rpm to produce either isobutanol or shikimate.

### Estimation of cell viability

Cell viability was estimated by plating the spent culture in LB and LB with Ampicillin (100ug/ml) plates. This is done to give a projection of the number of cells alive and the plasmid loss after various time points. Samples are sent for GC analysis/HPLC performed with the corresponding media standards. We estimate isobutanol and ethanol produced along with the remaining glucose concentration, as well as other metabolites like formate, lactate, acetate, succinate and pyruvate.

### HPLC analysis

For the detection of both Shikimate and Isobutanol,1ml of the culture was spun at 4000 g for 5 minutes, followed by further spinning of the supernatant at 14.8 g for 5 minutes. About 50ul of supernatant was analysed in HPLC. To perform the HPLC Aminex 87H column (Biorad) was used as stationary phase and 5mM H2SO4 was used as the mobile phase at 0.750ml/min flowrate with RID detector which is useful for detecting monosaccharides and organic acids at the same time.

## Results

Looking at a metabolic map it is might be a non-controversial conjecture, for the carbon flux to proceed towards the production of higher isobutanol or shikimate, lesser amount of nitrogen input is required. In fact, Fig 2 shows the number of steps from glucose to the production of isobutanol (15 steps) and shikimate (12 steps) which don’t have nitrogen. Also, BL21 strain was used for the production of isobutanol due to valine-feedback independent acetolactate synthase (ilvG) but the product yield for shikimate was also higher using the same strain and its knockouts (Supplementary Fig S1) for unknown reasons. Fig 3 shows the essential requirement of aromatic amino acids in order to maintain cell viability for shikimate production. Whereas, for isobutanol production this is not essential since there are no biochemical repercussions affecting cell viability. Hence, isobutanol production is an ideal candidate for testing the effect of limited nitrogen source on product yield. Fig 3 shows that the consumption of glucose remains same in both nitrogen and nitrogen depleted conditions however, the production of isobutanol is increased by almost 30% in media devoid of nitrogen (i.e. 0% N). But this cannot be directly applied for increasing shikimate production as aromatic amino acid synthesis is downstream to shikimate unlike valine.

**Figure 2:**
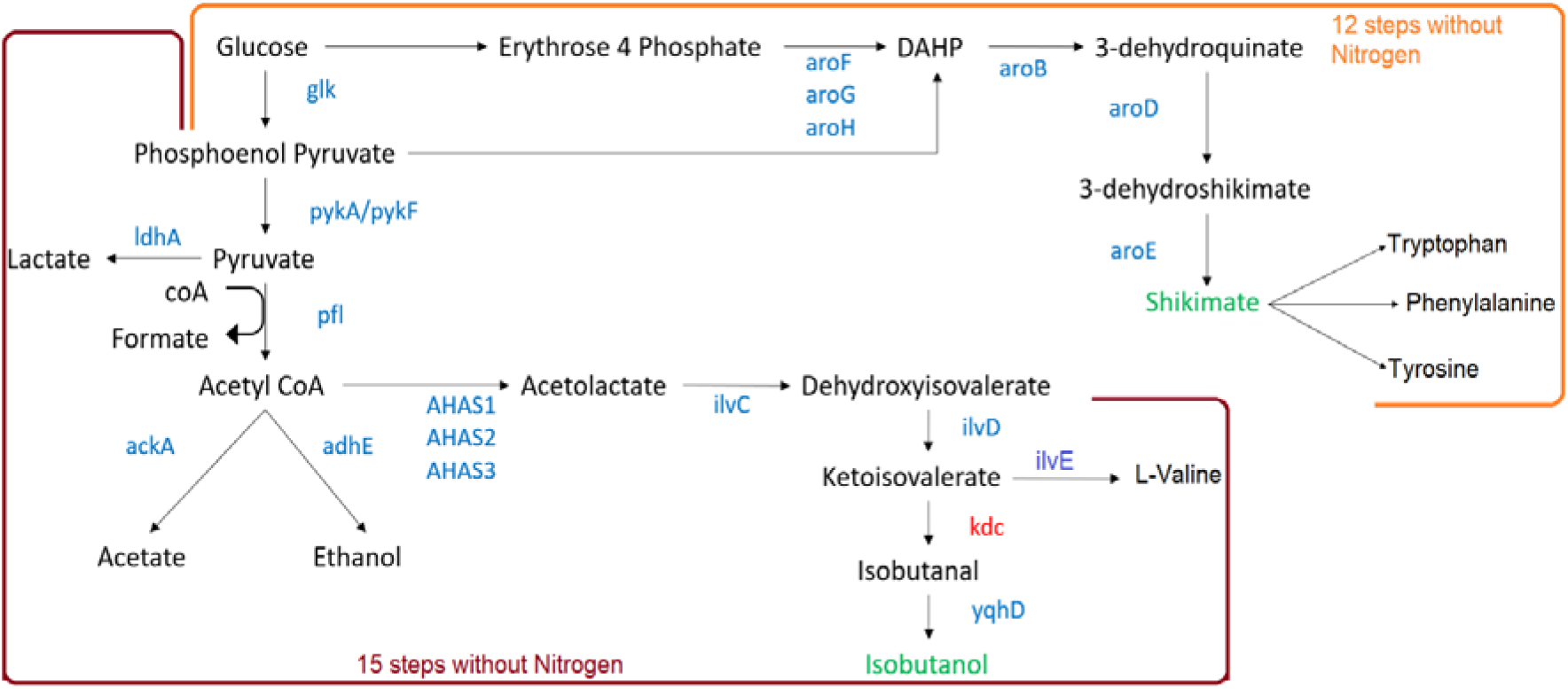
Pathway for Isobutanol and Shikimate production. The genes coloured blue is native to E. coli whereas the genes labelled in red are heterologously expressed and metabolites in green are products of interest.

**Figure 3:**
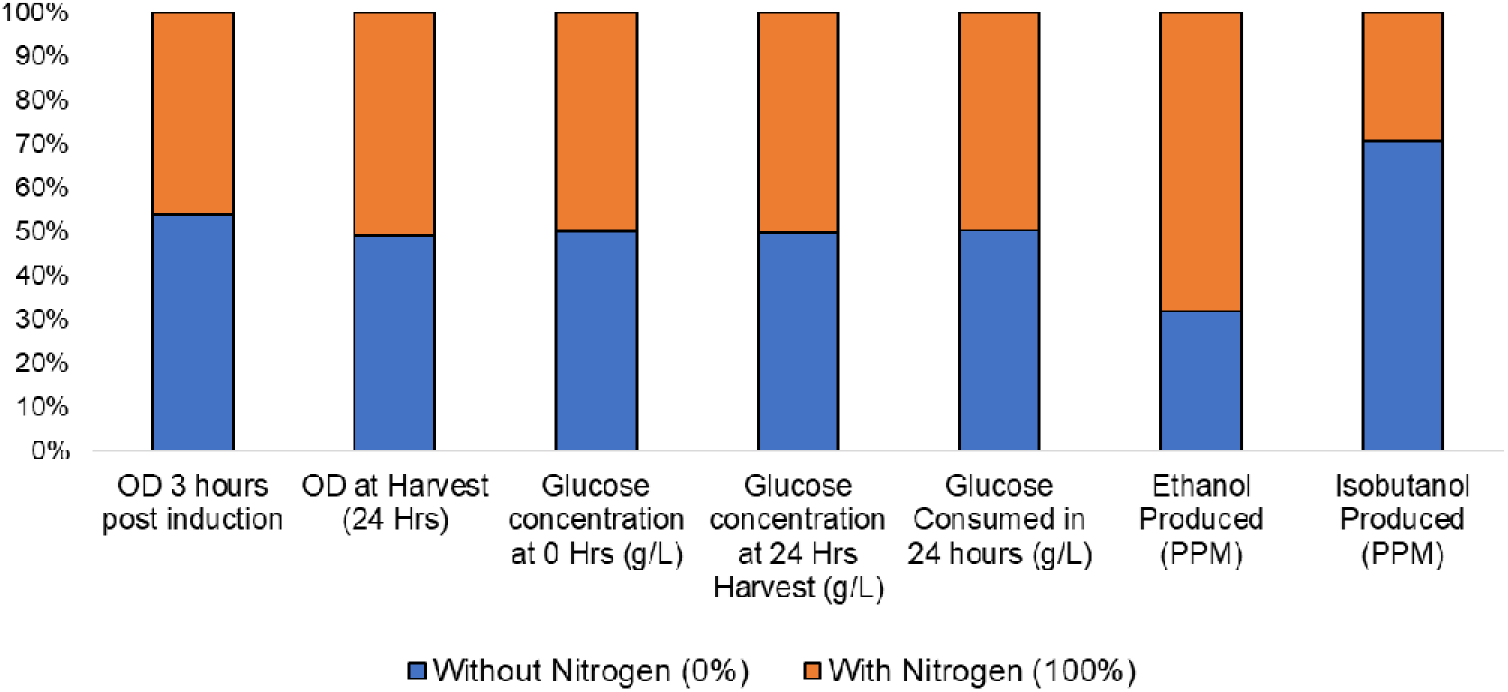
Graphical representation of initial shake flask experiment with the BL21 ΔadhE:ΔackAΔldhA with heterologous expression of KivD in media with and without nitrogen source.

The POSYBAL platform was used to identify the knockouts required for increasing the shikimate/isobutanol yield. Knockouts were derived by observing the flux through metabolite pathways. A negligible flux essentially represents a knockout. Hence, as depicted in Fig 1C a matrix of multiple solutions was used for finding the best fluxes for isobutanol and shikimate production. A set of 10^5^ iterations (Fig 4A) was used for simulating isobutanol and 10^4^ iterations (Fig 4B) were used for simulating shikimate production. Each dot represents one solution among the solution space. The optimal knockouts (encircled in Fig 4) for increased biomass against product yield for isobutanol is ΔackA: ΔldhA: ΔadhE triple knockout and shikimate yield is ΔaroA:ΔaroK:ΔaroL knockout. Further, to prove the significance of POSYBAL platform, about 10^4^ iterations were done for ΔackA, ΔldhA, ΔadhE in various combinations. This produces a scatter plot of bacterial population producing isobutanol. These ‘knockouts’ are produced by introducing negligible flux through the respective genes. Fig 5A shows the progression of population towards the ‘peak’ of the population distribution as the flux reduces from Wildtype (BW25113) to single, double and eventually triple knockout of ΔackA, ΔldhA, ΔadhE. Hence, it is easier to understand a population behaviour using this platform unlike ODE and FBA simulations. Fig 5B shows the normalized HPLC data for the wild type (BL21) and ΔackA:ΔldhA:ΔadhE triple knockout. There is an increased production of isobutanol in nitrogen depleted conditions especially at 3% nitrogen and a decreased production of acetate and no production of lactate in the triple knockout.

**Figure 4:**
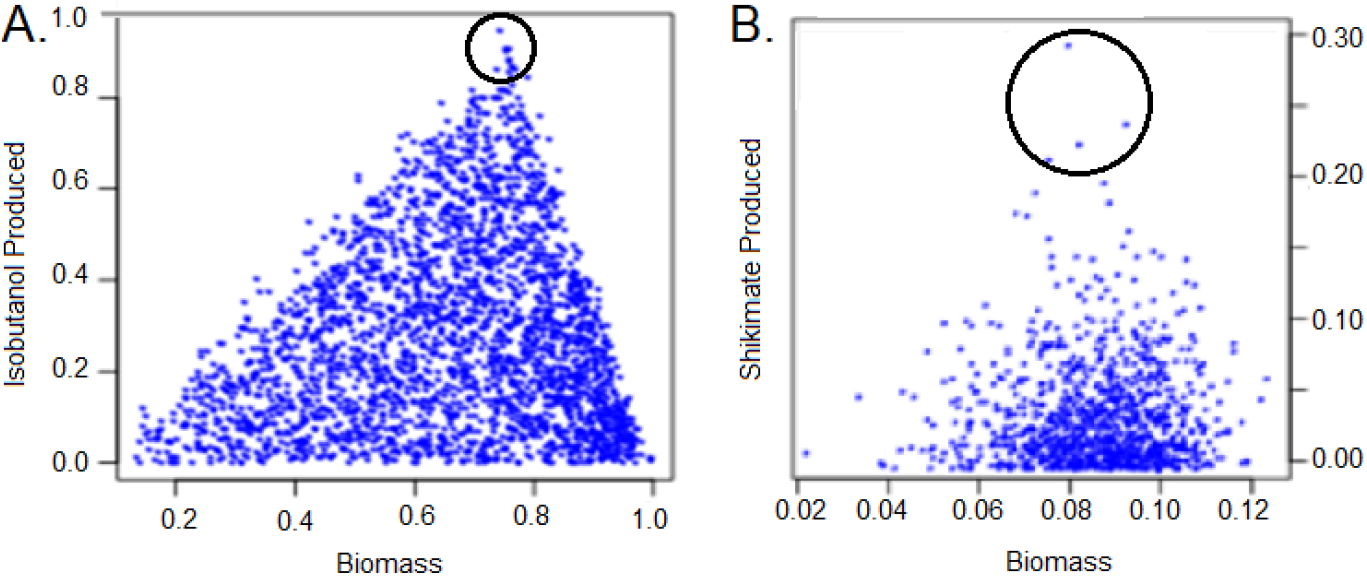
The figure represents the scatter plot of the population distribution for normalized isobutanol/shikimate production vs biomass A. Scatter plot of the population distribution for isobutanol production vs biomass comprising the triple knockouts adhE, ackA and ldhE with the threshold considered (encircled). B. Scatter plot for shikimate production comprising of the triple knockout aroK, aroA and aroL which are one of the solutions (encircled) in the plot.

**Figure 5:**
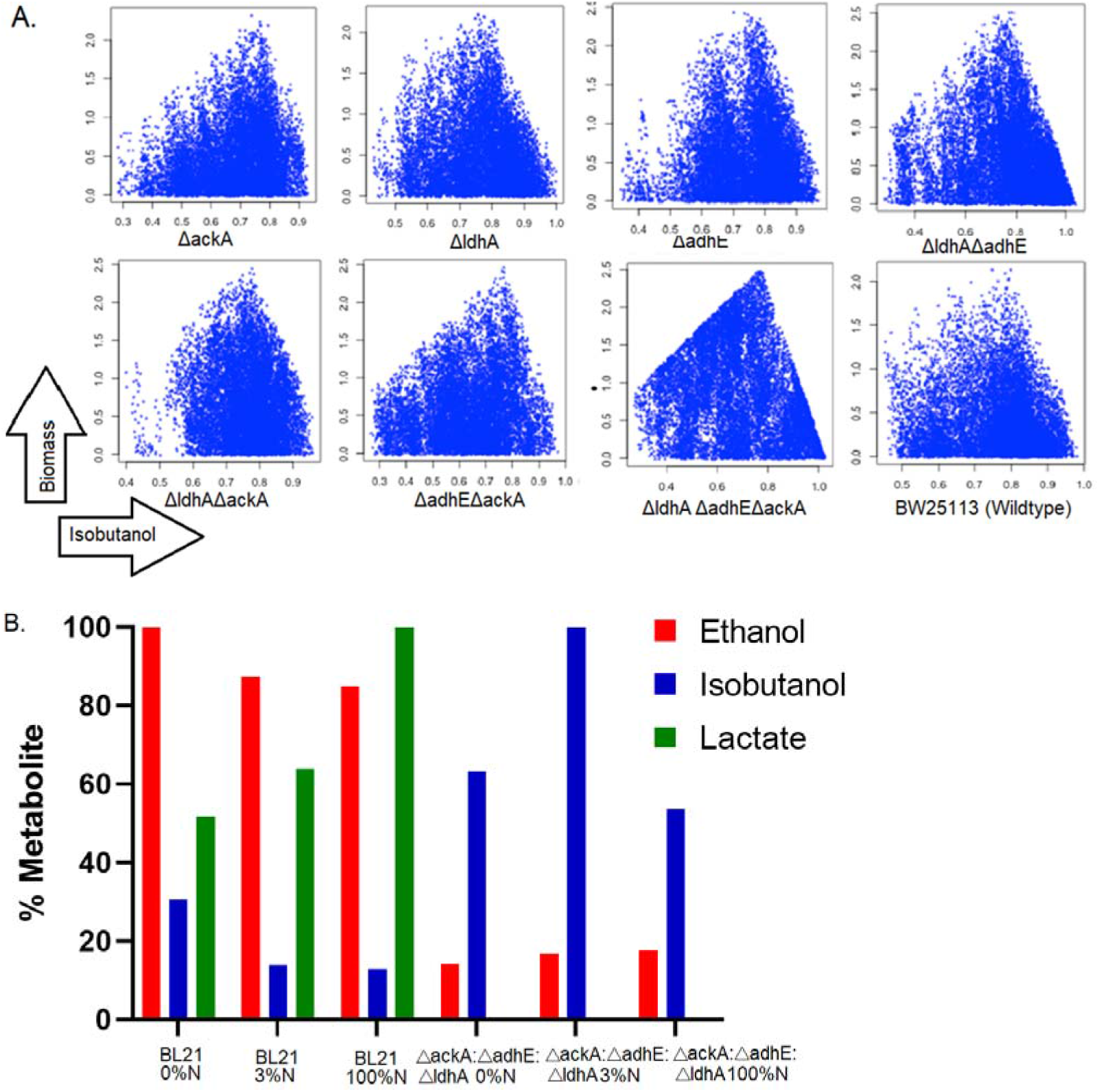
POSYBAL simulations for isobutanol production with the predicted knockouts. A. The population of bacterial cells producing isobutanol increases as flux decreases across adhE, ackA and ldhA genes. B. Graph depicting the normalized HPLC peaks for understanding the flux of Ethanol, acetate, lactate and Isobutanol in Wild type BL21 and ΔackA: ΔadhE:ΔldhA triple knockout expressing KivD. There is considerable reduction of flux towards acetate, lactate and acetate production compared to isobutanol.

For shikimate production the minimal requirements are indirect as the aromatic amino acid (essential) production is downstream. It is observed that despite intuitively choosing ackA based on Fig 2 as a knockout for producing more shikimate, the in vitro output proves otherwise (Fig 6B). POSYBAL simulations were performed for some of the knockouts derived from Fig4B. The individual scatter plots (Fig 6A) show inverse triangle relationship for aroL, pykA/F and ptsG. Hence, in these cases knockouts are important for increasing product yield. Whereas, gaussian plots are seen for ackA and partially for poxB. The scatter plots for knockouts which showed higher shikimate production. The shake flask experiments were performed with the previously predicted ΔaroK:ΔaroA:ΔaroL knockout along with ΔpoxB:ΔaroK:ΔaroA:ΔaroL and ΔptsG:ΔaroK:ΔaroA:ΔaroL quadruple knockouts (Fig 6B). The experiments were done with various nitrogen source (LB) concentrations of which the 20% LB (LB/5) was found to be the optimal concentration for maximum product yield. It is also seen that there is negligible flux of glucose without the nitrogen source. This shows that a certain amount of nitrogen is required for driving the carbon flux towards shikimate production. Fig 6B also shows the limited acetate required for higher shikimate yield in ΔaroK:ΔaroA:ΔaroL. The addition of poxB and ptsG knockouts reduce the acetate production by 68% (see supplementary table 2). This in turn increases the shikimate production in 20% LB (Fig 6B). Although the quadruple knockouts show that POSYBAL simulations help in understanding the population behaviour and flux of carbon, we sought the best triple knockouts which gave higher shikimate yield with enzymes such as tkaA and pntAB which showed direct correlation with shikimate production. These enzymes were taken from the ASKA collection and expressed in BL21 ΔaroK:ΔaroA:ΔaroL knockouts. The experiment was performed in bioreactor with 20% LB with 08 and 1.6% glucose. The production of shikimate after 24 hours in ΔaroK:ΔaroA:ΔaroL was 1634 ppm which was doubled by tktA overexpression to 3022 ppm and whereas, pntAB overexpression gave initial higher yields i.e. 1130 ppm (at 12 hours) but they remained in similar concentration (1245 ppm) after 24 hours as well (Fig 7A). For isobutanol production a fairly straightforward correlation was observed in 3% nitrogen containing complete minimal M9 media where the isobutanol concentration produced was 2235 ppm after 24 hours (Fig 7B).

**Figure 6:**
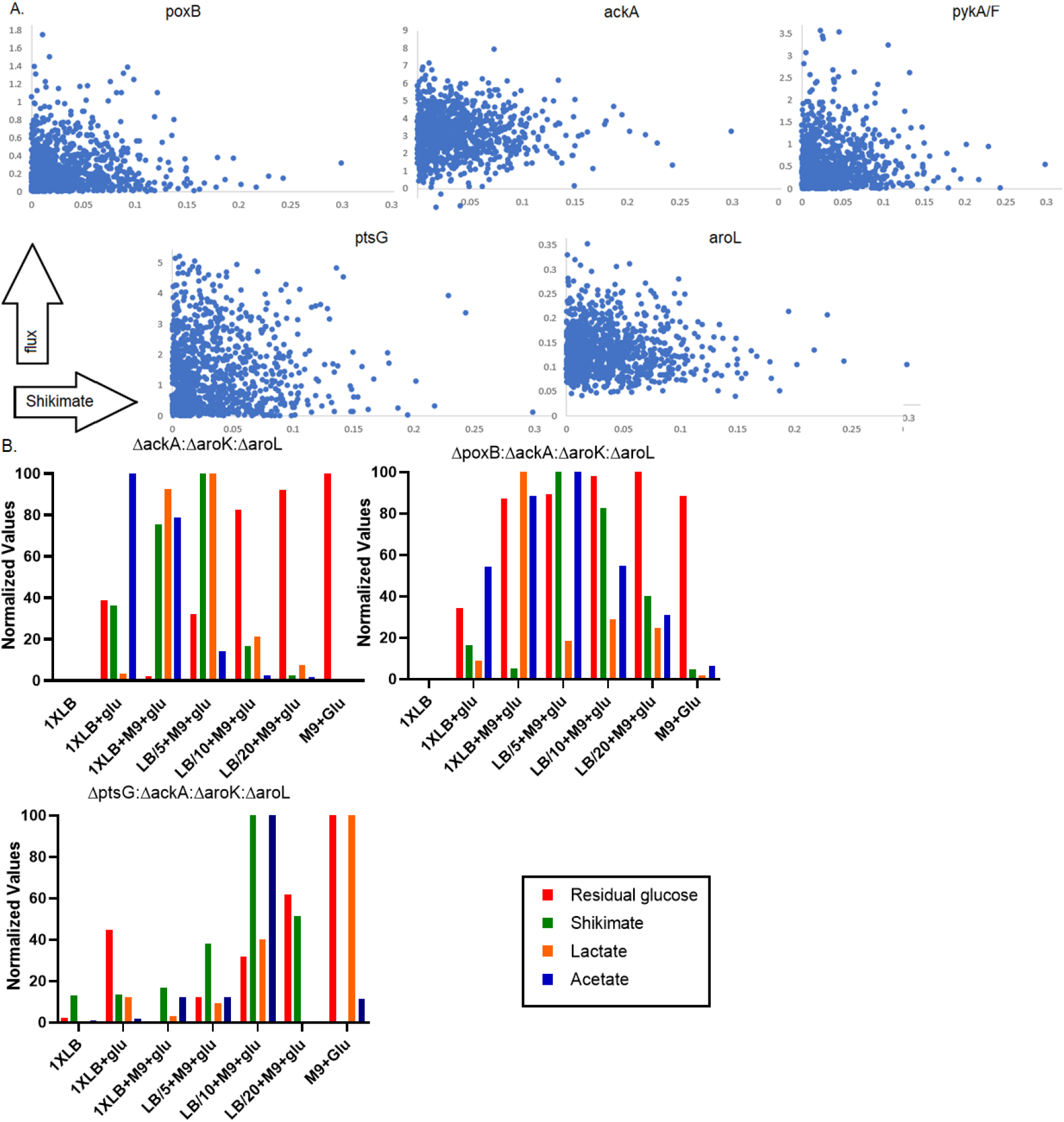
POSYBAL simulation for minimal requirements necessary in shikimate production. A: Scatter plot representing fluxes through each gene and the corresponding shikimate produced. It is observed that despite intuitively choosing ackA as a knockout for producing more shikimate, but the in vitro output proved otherwise. Through POSYBAL platform it is observed that poxB, pykA/F and aroL can be knocked out to produce high amount of shikimate whereas an ‘intermediate’ flux through ackA and ptsG produces higher shikimate than its knockout. B: Normalized invitro graphs show that acetate production is required for the flux to move towards shikimate production. Also, glucose cannot be consumed when a nitrogen source (LB) isn’t available.

**Figure 7:**
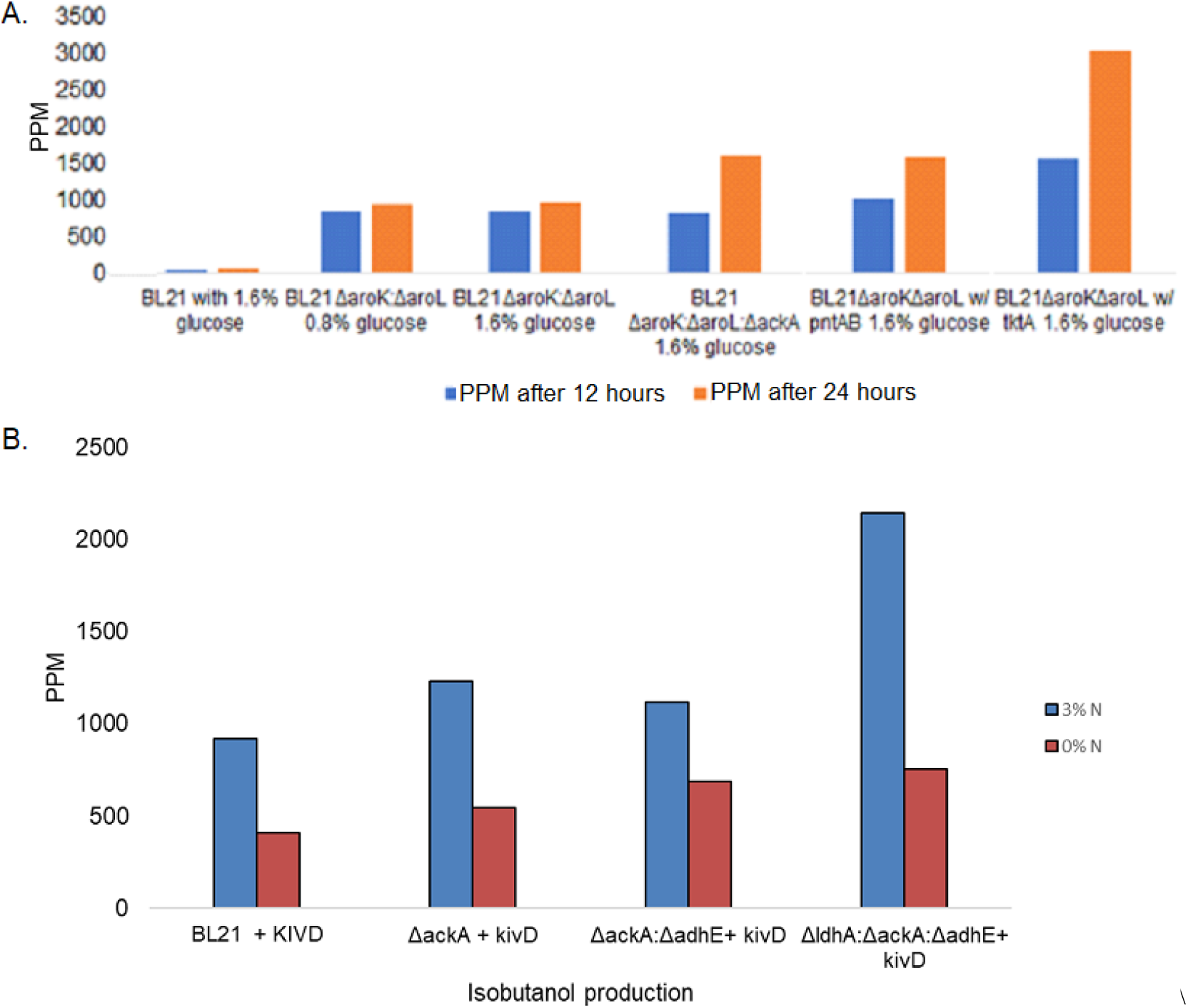
A. Production of Shikimate (mg/L or PPM) in BL21, ΔaroK:ΔaroL, ΔaroA:ΔaroK:ΔaroL, ΔaroA:ΔaroK:ΔaroL with pntAB and tktA overexpression in LB/5 or 20% LB (nitrogen source) in bioreactor. B. Production of Isobutanol in 0% and 3% nitrogen after 24 hours in pilot scale fermenter.

## Discussion

Traditionally, the production of butanol was done by ABE (Acetone, n-butanol and Ethanol) fermentation method. The carbon source used during this anaerobic fermentation procedure was starch with Clostridium acetobutylicum (or C. beijerinkii) as a whole-cell catalyst (Kujawska *et al*, 2015). The solvents were produced in a ratio of 3 parts Acetone, 6 parts Butanol and 1-part Ethanol. In the post genomic era, the use of systems biology and heterologous gene expression have ushered a revival in “bio-butanol”. (Atsumi et al. 2008) showed that the integration of the Ehrlich pathway into the branched chain amino acid pathway was sufficient to generate isobutanol in *E. coli* under non-fermentative conditions. Also, the addition of the heterologous gene kivD (ketoisovalerate decarboxylase) from *L. lactis* is required to produce isobutanal which is converted to isobutanol by multiple native isobutyraldehyde dehydrogenases such as YqhD, AdhP, FucO, EutG, YaiY, BetA, EutE and YjbB (Rodriguez and Atsumi, 2012) (Fig 2). Historically, Shikimate is produced by using plant sources, since they contain similar biosynthetic pathways. Star anise (*Illicium anisatum*) is usually used for extracting shikimate (1.5% w/v) (Bradley M, 2005). A better substitute is Sweetgum (*Liquidambar styraciflua*) which has a product yield of 2.4-3.7% w/v (Enrich *et al*, 2008). In *E. coli* It is obtained by knocking out genes such as aroK, aroL which participate in aromatic amino acid biosynthesis (Chen *et al*, 2012). Engineering microorganisms is imperative for higher yields or easier extraction processes.

However, static biochemical maps often don’t portray the nuances of dynamicity in aa cell. Even, traditional FBA or ODE models would give a single solution which may or may not work in vitro. However, POSYBAL simulations are different from traditional FBA models as they consider the overall ‘presence’ of a metabolite ‘through’ a population of cells. Hence, multiple combinations entail this concept and result in a normal distribution of substrate vs metabolite. Although, this stochastic combination requires intensive computational power, it is useful for determining combinations which may not be always ‘intuitive’. The platform shows different phenotypes of a single strain in its population. Notice, that the ‘cells’ in the scatter plot (Randomly picked points from the solution space) portrays a specific behaviour of a particular phenotype’s reaction flux and its corresponding effect on the target metabolite of interest. In general, three types of patterns are observed namely, inverse triangle correlation where a knockout may result in the production of a target metabolite. A direct triangle shows that a gene overexpression is required for metabolite production. A random scatter means there is no correlation involved. In the current study we demonstrate this with two examples, the production of isobutanol is straightforward since, valine biosynthesis can continue with limited nitrogen and the carbon flux can be diverted towards the production of acetolactate (and subsequently isobutanol) through microaerophilic conditions. However, in case of shikimate this approach fails as a stoichiometric deficit is observed. An example of this is demonstrated here for the role of ackA in shikimate synthesis. One may assume that knocking out acetate production (ackA) in Fig 2 would divert the flux towards shikimate production. However, this is counter-productive as seen in Fig 9.

Intuitively it can be assumed that knocking out acetate production (ackA) in Fig 2 would divert the carbon flux towards shikimate production. However, this is counterintuitive as seen in Fig 9A. With FBA models a stoichiometric assumption can be made since 2 molecules of glucose (C_6_H_12_O_6_) are needed to produce one molecule of shikimate (C_7_H_10_O_5_). But, the atomic excess of C_5_H_14_O_7_ has to be converted or excreted in other metabolic forms else, the reaction can reverse in order to maintain the cell viability. With the POSYBAL platform it is feasible to find out if a knockout, knockdown or overexpression is required for optimal production of shikimate (or any other metabolite). It is observed that despite intuitively choosing ackA as a knockout for producing more shikimate, but the in vitro output shows otherwise. Through POSYBAL platform it is seen that poxB, pykA/F and aroL can be knocked out to produce high amount of shikimate whereas an ‘intermediate’ flux through ackA and ptsG (Fig 9A) produces higher shikimate than its knockout. Fig 9B shows that limited amount of acetate (ackA expression) is required for the production of Shikimate. When it comes to glucose media the acetate is upstream in production and subsequently lactate is also increased. But in case of LB media the input is predominantly consists of nitrogen sources and hence the carbon ‘system’ in turn is downstream. The enzymatic journey taken for production of acetate is lesser in case of glucose medium than LB media. Along with this principle, it is important to consider the limiting effects of nitrogen for production of non-nitrogenous metabolites. This, along with an understanding of the anaerobic *E. coli* biochemistry, helps us in devising a strategy of N-swap coupled with partial aerobic flow through the fermentation process to maximize isobutanol/shikimate production. It is also important to note that the POSYBAL simulations were done with BW2511 strain whereas, with the exception of initial experimentation, the BL21 strain was used for producing shikimate and isobutanol. However, these strains have near identical genes and pathways and hence, the simulations stick to the in vitro repertoire.

The results from the insilico simulations were tested in the lab by creating a triple knockout ΔackA:ΔadhE:ΔldhA using p1 transduction method. Additionally, ketoisovalerate decarboxylase (kivD) was introduced to convert ortho-isovalerate to isobutanal. Minimal media was utilized to convert glucose (carbon source) to isobutanol in shake flasks. Initial experiments showed higher titre of Isobutanol yield in media devoid of Nitrogen (Fig 4). Whereas, in Shikimate, ΔaroA:ΔaroK:ΔaroL knockouts fail to survive as an external source of aromatic amino acids is required for continued survival of the cells.

The validation of computer predictions showed an increased flux of isobutanol away from ethanol production as seen in Fig 7 and 8. the bioconversions and knockouts were done in BL21 strain. The overall yield of shikimate increased from ~50 to 100 ppm in BL21 strain compared to the BW25113 (See supplementary Fig S2) also, aroK and aroL knockouts of BW and BL21 showed the same ‘signature’ of higher yield and BL21 was used to make ΔaroA:ΔaroK:ΔaroL triple knockout. The optimal biomass to isobutanol reduction ratio is observed in 3% nitrogen in minimal media. Similarly, higher yield of shikimate was seen in the predicted ΔaroA:ΔaroK:ΔaroL triple knockout compared to the double knockout (ΔaroK:ΔaroL) and wild type (Fig 9).

Based on the threshold parameters as mentioned and selected earlier by POSYBAL simulations, it was found that the triple knockout of ΔadhE:ΔackA:ΔldhA triple knockout corresponds to 2770 solutions of the 100000 (Fig 5a) solutions obtained, which is basically 2.7% of the solution set. Similarly, for shikimate 300 solutions were obtained of which five solutions (Fig 5b) were seen as optimal (16%). We see that there is no central governing systemic intelligence to a collection of the reaction set that has a small section (probability) wherein the system produces the metabolite of interest (in this case isobutanol) and this probability increases when the stem is reengineered (in case of shikimate) wherein certain gene expressions are blocked (knocked out). A filter to locate solutions that have reduced flux through specific reactions below a threshold finally points to an optimal knockout.

## Conclusion

The in-silico platform for various species of bacteria like *E. coli, M. tuberculosis, P. aeruginosa, C. acetobutylicum* have been described previously. They are either based on the non-linearity of interconnected ordinary differential equations (ODE) that represent various enzymes which describe the cellular interactions with its intrinsic kinetic parameters or in a Flux based mode where stoichiometry of each metabolite is linearly connected to another of the ensemble. Both these in silico modes essentially describes the functionality of a single cell and assumes homogeneity of behaviour in a population of an isogenic bacteria. In reality, a population of bacteria is not only asynchronous in its physiological state, no two cells in a population are in metabolic congruence. In conventional FBA the optimal solutions derived out of maximization/minimization of a particular reaction gives an understanding of the system required to achieve a theoretical maximum/minimum in a utopian environment. A living cell can be visualized as an ensemble of underdetermined equations that connects metabolites which produces an infinite array of possible solutions. Each cell in the population may use one set of these solutions. This tacitly entails that no two cells have identical expression levels in a given environment. These varying behavioural signatures enables the system to be robust enough to handle stress factors such as nutritional deficiency, osmotic imbalance, temperature shock or presence of antibiotics. These kind of competitive growth “advantage-disadvantage” simulation can be generated in our POSYBAL population model. It is seen that even in media with optimum nutrient availability cells diverge in their growth rate and rapidly move away from being synchronous. This divergence that is seen experimentally and is a natural outcome of our POSYBAL platform. It is also seen that this divergence of synchronicity is dependent on the flux through some key non-essential pathways wherein specific knockouts produce altered divergence.

## Acknowledgement

The financial support from BIG grant (Grant no. BIRAC/CCAMP0186/BIG-04/14) to Janani Venkataraman and DBT BBRC for RICEFUEL (Grant no. BT/IN/Indo-UK/SuBB/21SSY/2013) grant to Ramanujan K S and Santanu Datta is gratefully acknowledged. We acknowledge the initial computational work done by Madhura Adavkar, Vetri Selvi and experimental work by Rajeswari Basu.

## Author contribution

RKS drove the initial formulation and validation of the population model. SR programmed the in-silico platform and drove the entire in-silico cross validation with RKS. DM and RVH validated the platform by performing the experiments for isobutanol and shikimate production respectively. AG performed the scaled-up version of the experimental procedures using bioreactor. APS assisted SR and DM in performing experiments and was one of the major contributors in pooling the diverse experimental data into a manuscript format. NK supervised the making of the multiple constructs and directed the experimental platform for isobutanol production while JV oversaw the shikimate project. SD conceived the idea for population model and nitrogen swap in media.

