## Supplementary for "Maximization of non-nitrogenous metabolite production in *E. coli* using population systems biology"

**Supplementary methods**

1. **Preparation of E. coli knockouts**:

Making the P1 lysate:

A 5 ml culture of the donor strain is grown at 37^o^C in LB medium with 0.2% glucose and 5 mM CaCl2 till it becomes barely turbid. 100μl P1 virulent (phage titer 109-1010) is added and continued to be incubated at 37◦C with good aeration until lysis occurs. After centrifugation the lysate is filtered out using a 0.45uM filter and stored at 4^o^C leaving behind the debris.

Performing the P1 transduction:

The recipient strain is grown overnight. 5ml culture is centrifuged at minimum g and resuspended in about 2.5ml of P1 solution (10mM CaCl2 + 5mM MgSO4).100μl of these cells are mixed in P1 with 1, 10, and 100 μl of phage lysate in different tubes. A control tube without phage lysate is also included. All the tubes are incubated for 20 minutes at 37^o^C. In order to stop further phage binding 200μl of 1M Na-citrate followed by 1ml LB are added to the tubes and incubated for 1hour at 37^o^C. Post centrifugation the cells are resuspended in 100μl LB and plated on selective plates containing 5mM Na-Citrate. The following day single colonies are streaked out on fresh selective plates with 5mM Na-Citrate. The transductants with the new gene fragment is confirmed by PCR.

Generation of multiple gene knockouts in BL21 strain for enhanced product formation:

In order to drive the metabolic flux towards the desired product formation systematic knockouts are made by utilizing the BW single knockout library. The gene is replaced with kanamycin cassette as marker, and to generate the next knockout the cassette is flipped out with pCP20 plasmid. The prime focus is to generate multiple gene knockout BL21 strain in order to prevent the formation of undesirable products and drive the flux towards the formation of desired product. Currently a triple knockout has been produced by re

moving ackA, adhE and ldhA genes preventing the synthesis of acetyl phosphate, acetate and lactate from pyruvate and the production of ethanol from acetaldehyde thus abating ethanol formation.

2. **Programming POSYBAL for Isobutanol**

\# POSYBAL_multiore.R

POSYBAL_multicore.R is a script that has been built to read an FBA model and parse into the linear inverse model. Further a single optimum solution is generated (Biomass maximisation) and used as a starting point for the population run. The model sparce matrix along with a starting point is used and it is parallelised into 6 cores to develop independently random population of x numbers. The individual files are populated further to form a mega collection of iteration that shall constitute the targeted population number. (Multiple different maximasation can also be generated, where each of the cores can start working with different optimum solution further giving much more diverse solution)

\## Installation

This script requires the user to install R programming language.

The following libraries are essential for running the script post the installation of R.

1. LIM library and its associated dependency

2. For each library and its associated dependencies

3. doMC library and its associated dependencies.

The model file must be present in the same directory as the script as the same is passed as an argument to the script.

\## USAGE

Rscript POSYBAL_multicore.R Ecoli_isobutanol.lim

\## Input

Ecoli_isobutanol.lim

\## Output

Ecoli_isobutanol_posybal_all.csv

\# ISBANAL.R

Once the Matrix of all the solution points containing different possible solution of an underdetermined system is generated, it is passed as an argument for this script.For the metabolite of interest the script filters out all the samples in the population having flux greater than the threshold selected (ex. 90% of max flux and above) generating the maxflux containing subset of the population as a file output. Further it also analyses in the respective population the fluxes of the reaction that are found to run minimal(10% of respective reaction max) allowing us to visualise the reaction that can be knocked out to harness maximum target metabolite reaction.

The final output is the list of reactions that are found to be running low from the given model for the thresholds selected and subset of the population corresponding to maximum target flux reaction.

\## USAGE

Rscript ISBANAL.R Ecoli_isobutanol_posybal_all.csv

\## Input

Ecoli_isobutanol_posybal_all.csv

\## Output

kncdwn.csv

maxisobutanol.csv

reactions_isobutanol.txt

\# extracting_selected.py

This script takes in the reported reaction list as the input file and pulls out the data (reaction equation, genese involved) from the model reaction only file(main.txt). The final output is an text file containing the reactions that are running low for maximum target metabolite flux with the details of the genes involved in the respective fluxes. These gene list can further be used to carry out knock outs

\## Installation

This script requires the user to install Python programming language

\## Usage

python extracting_selected.py

\## Input

main.txt

reactions_isobutanol.txt

\## Output

output_isobutanol.txt

3. **Programming POSYBAL for Shikimate**

\# POSYBAL_multiore.R
POSYBAL_multicore.R is a script that has been built to read an FBA model and parse into the linear inverse model. Further a single optimum solution is generated (Biomass maximisation) and used as a starting point for the population run. The model sparce matrix along with a starting point is used and it is parallelised into 6 cores to develop independently random population of x numbers. The individual files are populated further to form a mega collection of iteration that shall constitute the targeted population number. (Multiple different maximasation can also be generated, where each of the cores can start working with different optimum solution further giving much more diverse solution)

\## Installation
This script requires the user to install R programming language.
The following libraries are essential for running the script post the installation of R.
1. LIM library and its associated dependancy
2. foreach library and its associated dependancies
3. doMC library and its associated dependancies.

The model file must be present in the same directory as the script as the same is passed as an argument to the script.

\## USAGE
Rscript POSYBAL_multicore.R Final_ecoli_model.lim

\## Input
Final_ecoli_model.lim

\## Output
Final_ecoli_model_posybal_all.csv

\# ISSHIKIMATE.R
Once the Matrix of all the solution points containing different possible solution of an underdetermined system is generated, it is passed as an argument for this script.For the metabolite of interest the script filters out all the samples in the population having flux greater than the threshold selected (ex. 90% of max flux and above) generating the maxflux containing subset of the population as a file output. Further it also analyses in the respective population the fluxes of the reaction that are found to run minimal(10% of respective reaction max) allowing us to visualise the reaction that can be knocked out to harness maximum target metabolite reaction.
The final output is the list of reactions that are found to be running low from the given model for the thresholds selected and subset of the population corresponding to maximum target flux reaction.

\## USAGE
Rscript ISSHIKIMATE.R Final_ecoli_model_posybal_all.csv

\## Input
Final_ecoli_model_posybal_all.csv

\## Output
kncdwn.csv
maxshikimate.csv
reactions_shikimate.txt

\# extracting_selected.py
This script takes in the reported reaction list as the input file and pulls out the data (reaction equation, genese involved) from the model reaction only file(main.txt). The final output is an text file containing the reactions that are running low for maximum target metabolite flux with the details of the genes involved in the respective fluxes. These gene list can further be used to carry out knock outs

\## Installation
This script requires the user to install Python programming language

\## Usage
python extracting_selected.py

\## Input
main.txt
reactions_shikimate.txt

\## Output
output_shikimate.txt

**Supplementary Figures:**

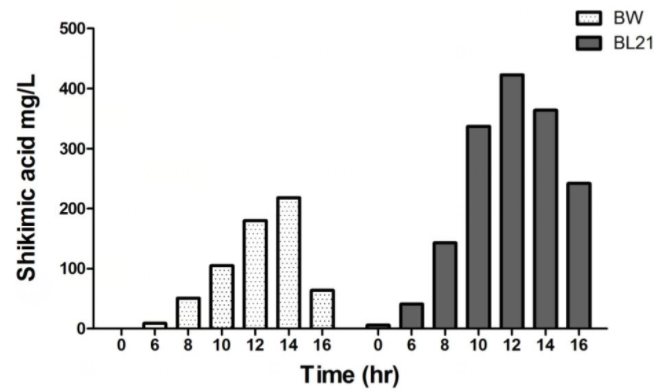

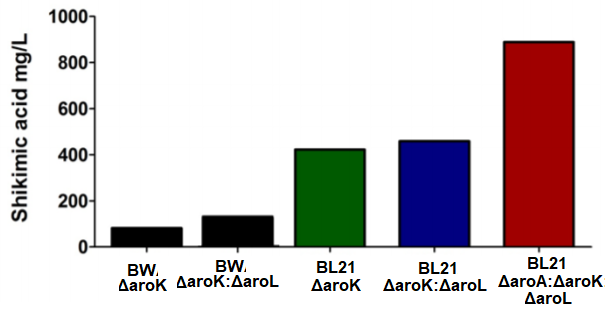

Figure S1: Comparison between BL21 and BW25113 strain for shikimate production done in 100 ml culture of M9 with LB. A. The overall yield of the production increased in the BL21 strain. B. The same phenomenon is seen in the ΔaroA:ΔaroK:ΔaroL knockouts in BL21 as compared to BW

Supplementary tables:

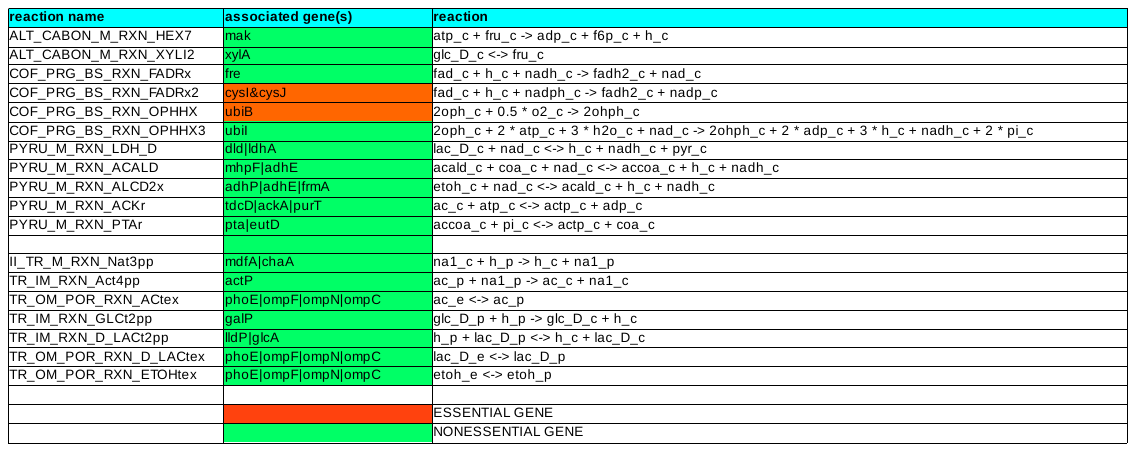

Supplementary Table 1 - The table below consists of the list of reactions that were found to be running less than 10% of the maximum corresponding to isobutanol value ranging from 80-100%. It entails the details of the reaction, the genes involved with it and the essentiality information. The genes highlighted in green are non-essential and can be knocked out in any combination or together whereas, the genes highlighted in red are essential and cannot be knocked out.

| ΔaroK:ΔaroA:ΔaroL |  |  |  |  |  |
| --- | --- | --- | --- | --- | --- |
|  | Final OD 600 | Residual glucose | Shikimate | Lactate | Acetate |
| 1XLB | 3.16 | 0 | 0 | 0 | 0 |
| 1XLB+glu | 4.15 | 2.96 | 23.75 | 69.14 | 1238.48 |
| 1XLB+M9+glu | 5.09 | 0.18 | 49.38 | 1948.33 | 974.75 |
| LB/5+M9+glu | 1.555 | 2.46 | 65.3 | 2107.19 | 175.95 |
| LB/10+M9+glu | 0.324 | 6.31 | 11.03 | 449.05 | 32.77 |
| LB/20+M9+glu | 0.105 | 7.05 | 1.76 | 163.55 | 19.69 |
| M9+Glu | 0.068 | 7.63 | 0 | 0 | 0 |
| ΔpoxB:ΔaroK:ΔaroA:ΔaroL |  |  |  |  |  |
|  | Final OD 600 | Residual glucose | Shikimate | Lactate | Acetate |
| 1XLB | 2.42 | 0.19 | 13.71 | 0 | 0 |
| 1XLB+glu | 3.68 | 2.81 | 50.87 | 211.06 | 277.47 |
| 1XLB+M9+glu | 5.19 | 6.8 | 25.57 | 2307.76 | 450.71 |
| LB/5+M9+glu | 3.41 | 6.96 | 241.32 | 432.6 | 508.9 |
| LB/10+M9+glu | 1.433 | 7.63 | 201.74 | 669.57 | 279.39 |
| LB/20+M9+glu | 1.005 | 7.78 | 104.77 | 567.89 | 157.69 |
| M9+Glu | 0.377 | 6.9 | 24.2 | 40.83 | 32.11 |
| ΔptsG:ΔaroK:ΔaroA:ΔaroL |  |  |  |  |  |
|  | Final OD 600 | Residual glucose | Shikimate | Lactate | Acetate |
| 1XLB | 2.7 | 0.204 | 3.76 | 0 | 4.1 |
| 1XLB+glu | 3.59 | 3.43 | 3.91 | 4.3 | 7.07 |
| 1XLB+M9+glu | 0.96 | 0.03 | 4.94 | 1.12 | 45.79 |
| LB/5+M9+glu | 0.394 | 0.96 | 11.08 | 3.22 | 45.9 |
| LB/10+M9+glu | 0.218 | 2.45 | 28.99 | 13.96 | 377.95 |
| LB/20+M9+glu | 0.116 | 4.72 | 14.93 | 0 | 0 |
| M9+Glu | 0.126 | 7.63 | 0 | 34.7 | 42.62 |

Supplementary Table 2 - The table below consists of the concentration of various metabolites in ΔptsG:ΔaroK:ΔaroA:ΔaroL, ΔpoxB:ΔaroK:ΔaroA:ΔaroL
